# Voltage-dependent inward currents in smooth muscle cells of skeletal muscle arterioles

**DOI:** 10.1101/211540

**Authors:** Alexandra V. Ulyanova, Roman E. Shirokov

## Abstract

Voltage-dependent inward currents responsible for the depolarizing phase of action potentials were characterized in smooth muscle cells of 4^th^ order arterioles in mouse skeletal muscle. Currents through L-type Ca^2+^ channels were expected to be dominant; however, action potentials were not eliminated in nominally Ca^2+^-free bathing solution or by addition of L-type Ca^2+^ channel blocker nifedipine (10 μM). Instead, Na^+^ channel blocker tetrodotoxin (TTX, 1 μM) reduced the maximal velocity of the upstroke at low, but not at normal (2 mM), Ca^2+^ in the bath. The magnitude of TTX-sensitive currents recorded with 140 mM Na^+^ was about 20 pA/pF. TTX-sensitive currents decreased five-fold when Ca^2+^ increased from 2 to 10 mM. The currents reduced three-fold in the presence of 10 mM caffeine, but remained unaltered by 1 mM of isobutylmethylxanthine (IBMX). In addition to L-type Ca^2+^ currents (15 pA/pF in 20 mM Ca^2+^), we also found Ca^2+^ currents that are resistant to 10 μM nifedipine (5 pA/pF in 20 mM Ca^2+^). Based on their biophysical properties, these Ca^2+^ currents are likely to be through voltage-gated T-type Ca^2+^ channels. Our results suggest that Na^+^ and at least two types (T- and L-) of Ca^2+^ voltage-gated channels contribute to depolarization of smooth muscle cells in skeletal muscle arterioles. Voltage-gated Na^+^ channels appear to be under a tight control by Ca^2+^ signaling.

## Introduction

Regulation of pressure and local blood flow occurs at the level of resistance arteries and arterioles. Under physiological conditions, resistance arteries and arterioles exist in the state of partial constriction, termed myogenic tone. Myogenic tone is considered to be an intrinsic property of arteriolar smooth muscle cells, which membranes depolarize in response to increase in the intraluminal pressure. In contrast to many other excitable tissues, oscillations of membrane potential and spike-like activity in arteriolar smooth muscles are mainly mediated by the activity of voltage-gated L-type Ca^2+^ channels [1]. These channels are open in response to the depolarization, provide Ca^2+^ influx into the smooth muscle cells, which in turn activates various voltage-gated and Ca^2+^-sensitive channels, and also initiates endothelial Ca^2+^ signaling to induce vasodilation [2]. Although a relationship between change in membrane potential and myogenic response is considered to be universal throughout various tissues, it may be regulated differently based on autoregulatory responses and channels expression [3–9].

Vascular resistance of skeletal muscle is a critical determinant of total peripheral resistance and blood flow as skeletal muscle receives about 20% of cardiac output at rest and up to 80% during exercise [9]. Despite the physiological importance of skeletal muscle vasculature, ionic mechanisms underlying physiological responses of arterial smooth muscle to depolarization are still poorly understood. Sponteneous action potentials have been previously recorded in skeletal muscle arterioles were [10–12], although their ionic basis have not been fully characterized. In arteries and arterioles, action potentials are thought to be independent of voltage-gated Na^+^ channels, as they could not be blocked by TTX [13, 14]. Nevertheless, Keatinge demonstrated that Na^+^ ions carry the depolarizing current during bursts of action potentials in strips of carotid artery [14, 15]. In addition, significant TTX-sensitive Na^+^ currents were found in smooth muscle cells isolated from rabbit main pulmonary artery [16, 17], human and rabbit aorta [18–20], and murine mesenteric and femoral arteries [21, 22].

We found that TTX-sensitive voltage-gated Na^+^ channels indeed contribute to depolarizing current in smooth muscle cells of skeletal muscle arterioles. Consistent with the findings of Keatinge [14, 15], currents through TTX-sensitive voltage-gated Na^+^ channels become apparent in nominally Ca^2+^-free solution. Our results also indicate that at physiological extracellular [Ca^2+^], these Na^+^ currents are suppressed by elevations in cytosolic [Ca^2+^]. Since specific blockers of voltage-gated Ca^2+^ channels suppress both the upstroke and the after-depolarizing components of action potentials, Ca^2+^ channels are thought to be the main pathway for the depolarizing current. Several studies have recorded currents through dihydropyridine-sensitive Ca^2+^ channels (Ca_V_1.2) in the arteries of various vascular beds [13, 23–26]. Therefore, it is generally accepted that Ca_V_1.2 is the principal sub-type of voltage-gated Ca^2+^ channels involved in excitation-contraction coupling of vascular smooth muscle cells [27]. In addition to the L-type, vascular smooth muscle cells also have T-type voltage-gated Ca^2+^ channels (for review, see: [28–30]). These channels were found in rat mesenteric arteries [24], rat and human cerebral arterioles [6, 13, 31], guinea-pig coronary artery [25], and human mesenteric arteries [13]. Notably, the mRNAs for T-type voltage-gated Ca^2+^ channel (Ca_V_3.1 and Ca_V_3.2) were detected in arterioles of skeletal muscle [32]. The role of T-type Ca^2+^ channels in smooth muscle of vasculature is not clear, as Ca^2+^ influx through Ca_v_3.2 T-type channels appears to be essential for relaxation of coronary arteries, probably by acting through activation of BK_Ca_ channels [33, 34].

We therefore electrophysiologically characterized arteriolar smooth muscle cells and the ionic currents underlying their exitability. More specifically, we developed a methodology that allows for the isolation and study of murine skeletal muscle arterioles in ex-vivo preparation. We described some of the voltage-dependent characteristic of the smooth muscle cells using voltage- and current-clamp recordings, including perforated whole-cell path-clamp technique. We confirmed that in addition to the previously recognized L-type voltage-gated Ca^2+^ channels, TTX-sensitive Na^+^ and T-type Ca^2+^ channels were present in our preparation. We examined potential contribution of these channels to voltage-dependent inward currents electrophysiologically and pharmacologically. Our data demonstrate two types of voltage-gated Ca^2+^ channels and TTX-sensitive Na^+^ channels contribute to the voltage-dependent inward currents in response to the depolarization in smooth muscle cells of skeletal muscle arterioles. Their physiological properties and hence relative contributions may change depending on the state of the arteiolar vasculature.

## Materials and methods

### Solutions

The dissection solution contained (in mM): 140 NaCl, 4 KCl, 1.5 MgCl_2_, 1 CaCl_2_, 10 HEPES, and 10 glucose. The extracellular solutions 1Ca, 2Ca, and 5Ca contained (in mM): 140 NaCl, 4 KCl, 1.5 MgCl_2_, 10 HEPES, 10 glucose, and corresponding concentration of CaCl_2_. The 0Ca solution contained 140 NaCl, 4 KCl, 4 MgCl_2_, 10 HEPES, and 10 glucose. Concentration of free calcium ([Ca^2+^_free_]) measured by Ca^2+^-selective electrode was 2-5 µM. The extracellular solutions 10Ca, 20Ca, and 20Ba contained (in mM): 150 NaCl, 10 HEPES, and corresponding concentration of either CaCl_2_ or BaCl_2_. The intracellular solution for whole-cell voltage-clamp experiments contained (in mM): 20 CsCl, 110 aspartic acid, 5 Mg-ATP, 4.5 CaCl_2_ ([Ca^2+^_free_] ≈ 100 nM), 10 HEPES, and 10 EGTA. The pipette solution for perforated patch-clamp experiments contained (in mM): 150 KCl and 10 HEPES. All solutions were at pH 7.3, 310-315 mosmole/kg and room temperature. All chemicals, unless specified, were purchased from Sigma, USA.

### Isolation of vessels

Arterioles with diameter of 21 ± 5 µm (n = 123) were enzymatically isolated from total of 42 mice (3-16 weeks old, weighing 15-28g, C57BL6 or C57BL10 genetic background) of either sex. Based on their diameter in the un-pressurized state and because the vessels were occasionally connected to muscle-less capillaries, these were the fourth-order arterioles. Animals were sacrificed by a cervical dislocation procedure. Semitendinosus and biceps femoris muscles were quickly excised from the animal and placed in the dissection solution. Muscles were then cleaned of fat and connective tissue, chopped into small pieces and transferred into 0.35% collagenase (Type 2, Worthington Biochemical, USA) dissolved in the same solution. The tissue was shaken at 130 rpm, 37C°, for 45-70 min depending on the animal’s weight. After digestion, the tissue was centrifuged for 45 sec at 1,000 rpm and gently washed with dissection solution containing 10% of fetal bovine serum to stop enzymatic reaction. 0.2 mL of digested tissue was added to the electrophysiology perfusion chamber mounted on an inverted microscope. An arteriole was gently pressed against the cover slip by a helping fire polished glass micropipette. The study conforms to the Guide for the Care and Use of Laboratory Animals published by the US National Institutes of Health (NIH Publication No. 85-23, revised 1996).

### Electrophysiology

The following protocol was used in order to ease formation of gramicidin-perforated patch-clamp configuration. First, gramicidin A was diluted in ethanol to a stock concentration of 10 µg/mL. In a separate light-protected tube, 1 µL of solubilizing detergent tetraethylene glycolmonooctyl ether was mixed with 1 mL of the pipette solution for perforated patch-clamp to facilitate grimicidin incorporation into the plasma membrane of the cells [35]. 20 µL of gramicidin A stock were added into the tube with detergent solution, and the mixture was gently rocked for about 30 min. Just before use, a sample of solubilized gramicidin was further diluted 4-5 times with the pipette solution for perforated patch-clamp. The patch-clamp pipette was dipped into the solution without gramicidin for approximately 10 seconds and then backfilled with the gramicidin-containing solution. Perforation was achieved within 10-15 min after formation of giga-seal contact. Axopatch 200B (Molecular Devices, USA) amplifier was used for voltage- and current-clamp experiments. Formation of giga-seal contact was monitored in the voltage-clamp mode. At first, 10 mV test pulses were applied from 0 mV holding potential. When the giga-seal was formed, the holding potential was switched to –70 mV. Capacitive transients recorded in response to the voltage step from –70 to –60 mV are illustrated in Fig 1A. Trace a illustrates capacitive transients measured with gramicidin-perforated patch clamp of smooth muscle cells that are situated in the vessel. The large peak current, slow time course of decay, and relatively large steady-state current showed that the apparent capacitance was large and the input resistance was low, as it is expected for electrically coupled cells [36]. In contrast to the results of Yamamoto et al. for mesenteric arterioles [37], we found that most of smooth muscle cells in skeletal muscle arterioles are coupled.

**Fig 1.**
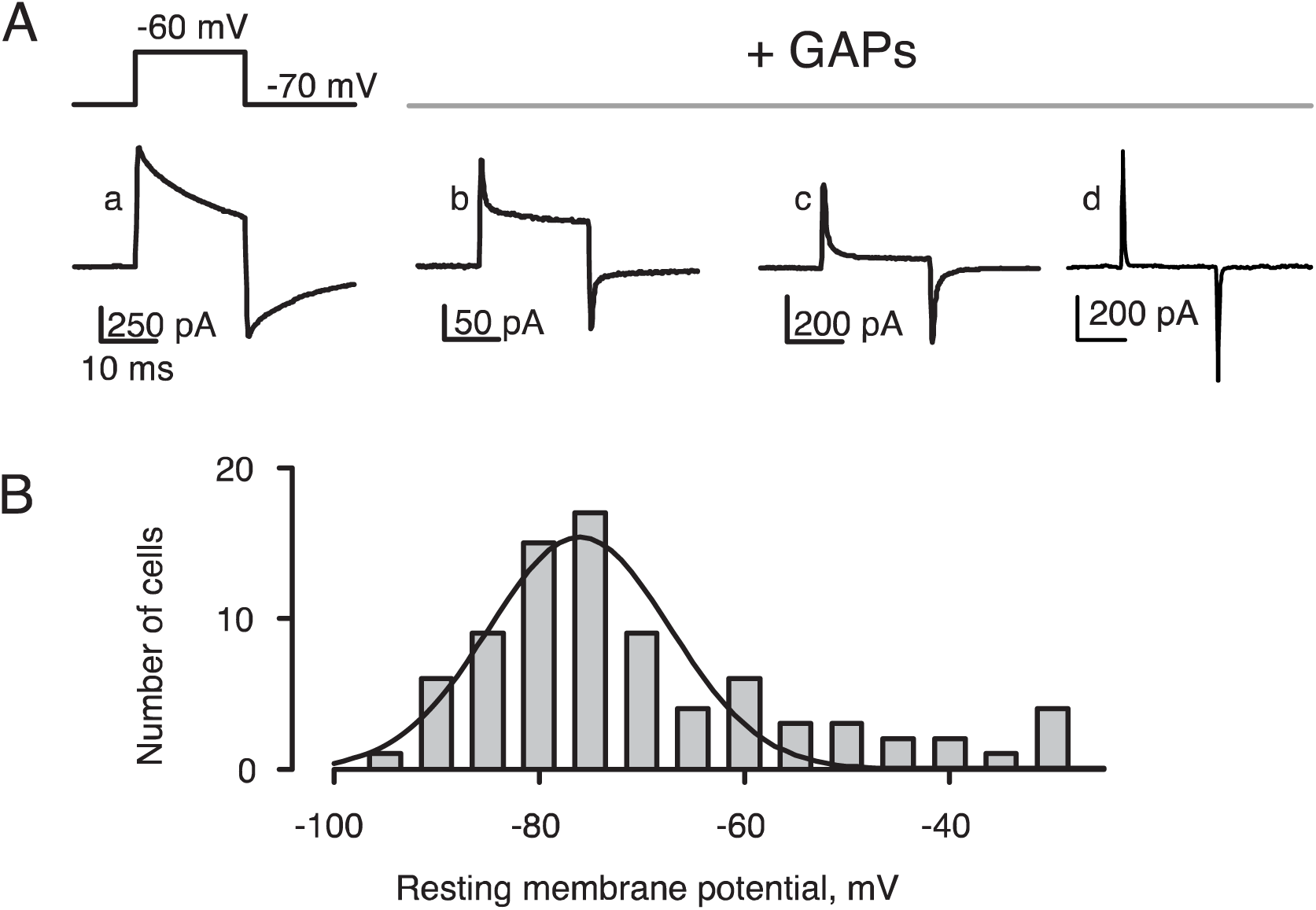
Electrical properties of smooth muscle cells. **A,** Capacitive transients were measured in response to the voltage step from –70 to –60 mV. The electrically coupled cells (trace a) became uncoupled after treatment with the gap-junction inhibitory peptides (GAPs, see Methods) during gramicidin-perforated patch clamp experiments (trace b-c) and after establishing the conventional whole-cell approach (trace d); **B,** Distribution of the resting membrane potential values was fitted by a single Gaussian function peaking at –77 ± 2 mV (smooth line). The average resting potential was V_rest_ = –68 ± 2 mV, n = 81.

To inhibit formation of gap junctions and thus electrically uncouple cells, tissue was incubated for 30 min in the dissection solution supplemented with 100 µM of ^37,43^GAP_27_, ^40^GAP_26_, and ^40^GAP_27_ peptides [38]. Input capacitance decreased and input resistance increased in vessels treated with the peptides (Fig 1A, traces b and c). In about two-thirds of cases, capacitive transients were like those in trace c. Input resistance, membrane capacitance, and series resistance were determined from capacitive transients [39]. Perforation by gramicidin was considered as satisfactory after the series resistance became less than 30 MΩ. Trace d in Fig 1A illustrates capacitive transient recorded after establishing the ruptured whole-cell approach in a vessel pretreated with gap-junction inhibiting peptides. Averaged membrane capacitance and input resistance of cells recorded with ruptured whole-cell approach were C_m_=14 ± 1 pF, n=42. Recordings were done in cells with apparent input resistance of at least 2 GΩ.

Voltages were recorded using gramicidin-perforated patch configuration and the fast current-clamp mode of the Axopatch 200B amplifier. The average resting potential was –68 ± 2 mV, n=81. Fig 1B shows the distribution of resting membrane potential values. The smooth line is the best fit by a single Gaussian function peaking at –77 ± 2 mV. Based on these observations, we analyzed only cells with the resting membrane potential more negative than –60 mV. Ionic currents were recorded using the ruptured whole-cell patch-clamp approach in cells with series resistance 3-10 MΩ. The holding potential for voltage-clamp experiments was –70 mV. Leak and capacitive currents were compensated electronically and by further subtraction of symmetric currents elicited with control pulses from –70 to –80 mV. Tracings were discarded if series resistance and/or membrane capacitance changed during the experiment by more than 5%.

### Statistics

Measured values are presented as sample average ± SEM. The statistical significance of the difference in averages was established using the two-tailed *t*-test (at the p < 0.05 level) using the normal distribution function and conventional estimates of standard deviation of the difference. In Fig. 2C, statistical significance of the difference of maximal rates of upstroke was established with the paired *t*-test. Curve fitting was done by a nonlinear least-squares routine of Sigma Plot (SPSS Inc., USA).

**Fig 2.**
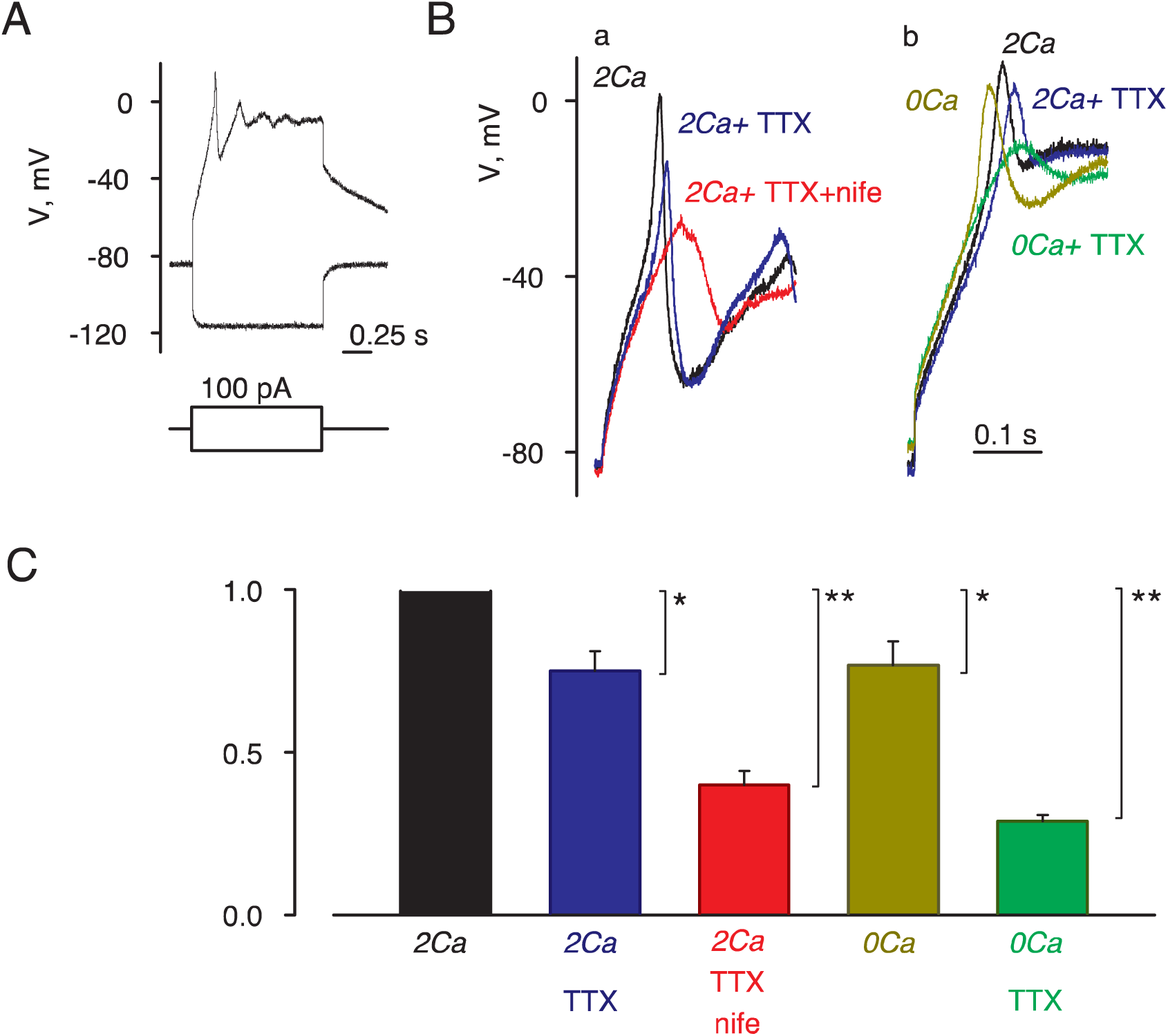
TTX affected action potentials in the absence of extracellular Ca^2+^. **A)** Hyperpolarizing current steps produced only passive voltage responses during gramicidin-perforated patch clamp experiments whereas depolarizing current steps induced action potentials starting from the threshold of about –50 mV. **B)** Effects of 1 µM TTX (blue trace) and 10 µM nifedipine (red trace) are shown in panel a at normal physiological conditions (2Ca solution); in nominally Ca^2+^-free (0Ca) solution, application of 1 µM TTX had much greater effect as shown in panel b. **C)** Averaged effects of TTX and nifedipine on the maximal rate of upstroke. Maximal rates of upstrokes recorded in different solutions were normalized to that recorded in 2Ca. TTX and nifedipine significantly reduced the maximal rate of upstroke in 2Ca solution. The effect of TTX was significantly greater in 0Ca than in 2Ca solutions. Asterisks indicate significance of the difference from control values in the *2Ca* solution (*, *P*<0.05; **, *P*<0.01).

## Results

### TTX affected action potentials in the absence of extracellular Ca_2+_

We characterized active responses to current injections in smooth muscle cells as shonw in Fig 2A in a representative current-clamp experiment. A 100 pA hyperpolarizing current step produced a passive response typical for an uncoupled cell (Fig 2A, bottom trace). Depolarizing current of the same strength evoked an active response that began with a rapid upstroke followed by series of depolarizing afterpotentials (Fig 2A, top trace). The initial upstroke started from the threshold of about –50 mV (Fig 2A). In contrast to the previous reports [14, 40], the upstroke was somewhat slower and of smaller amplitude in the presence of 1 µM of Na^+^ channel blocker TTX (Fig 2Ba, black and blue trace). In the bathing solution containing 1 µM of TTX and 10 µM of L-type Ca^2+^ channel blocker nifedipine, the upstroke was further reduced but not completely abolished (Fig 2Ba, red trace). Since the depolarizing inward current is thought to be primarily through Ca^2+^ channels, we tested how the upstroke is affected by the removal of extracellular Ca^2+^ (Fig 2Bb). The upstroke was not eliminated in nominally Ca^2+^-free (0Ca) extracellular solution, consistent with the idea that both Ca^2+^ and Na^+^ ions contribute to the depolarizing current. Importantly, TTX had much stronger effect in 0Ca, rather than in 2Ca, solution (Fig 2Bb, green and blue traces). The averaged effects of the channel blockers and Ca^2+^ removal on the maximal rate of the upstroke are compared in Fig 2C. Both TTX and nifedipine significantly reduced the maximal rate of the upstroke.

The results in Fig 2C have several implications about ionic mechanisms of action potentials in our experimental system. First, contributions of TTX- and nifedipine-sensitive components are not additive. The observation that TTX had much stronger effect at low, rather than at normal Ca^2+^, indicates that TTX-sensitive Na^+^ channels were inhibited, or not accessible for TTX, at normal Ca^2+^. Second, a part of depolarizing current appears to be resistant to both TTX and nifedipine. Therefore, we undertook the following experiments to investigate depolarizing ionic currents and whether or not voltage-gated Na^+^ channels are suppressed during the upstroke at physiologically normal [Ca^2+^].

### TTX-sensitive Na^+^ currents

To dissect inward ionic currents, we used the ruptured whole-cell voltage-clamp with cesium-based pipette solution designed to block outward K^+^ currents. Ionic currents evoked by steps to different voltages in 2Ca solution are illustrated in Fig 3A (left panel). The thick line highlights currents at 10 mV. It shows that voltage-dependent inward current had at least two kinetically distinct components. The more rapid component was through voltage-gated Na^+^ channels, as it was completely blocked by application of 1 µM TTX (middle panel). The slower current was likely to be through Ca^2+^ channels, as its magnitude increased with [Ca^2+^] (right panel). Averaged peak current–voltage relationships are shown in Fig 3B. The maximal density of TTX-sensitive current calculated as the difference between the maximal current densities before and after application of TTX was 20 ± 2 A/F with 2 mM Ca^2+^ in the bath. In the absence of TTX, peak current amplitudes declined with [Ca^2+^]. However, peak current amplitudes recorded in the presence of TTX increased with [Ca^2+^], indicating that the TTX-resistant current was entirely through Ca^2+^-selective channels. The maximal density of TTX-sensitive current dropped to 4 ± 1 A/F when extracellular Ca^2+^ was changed from 2 to 10 mM.

**Fig 3.**
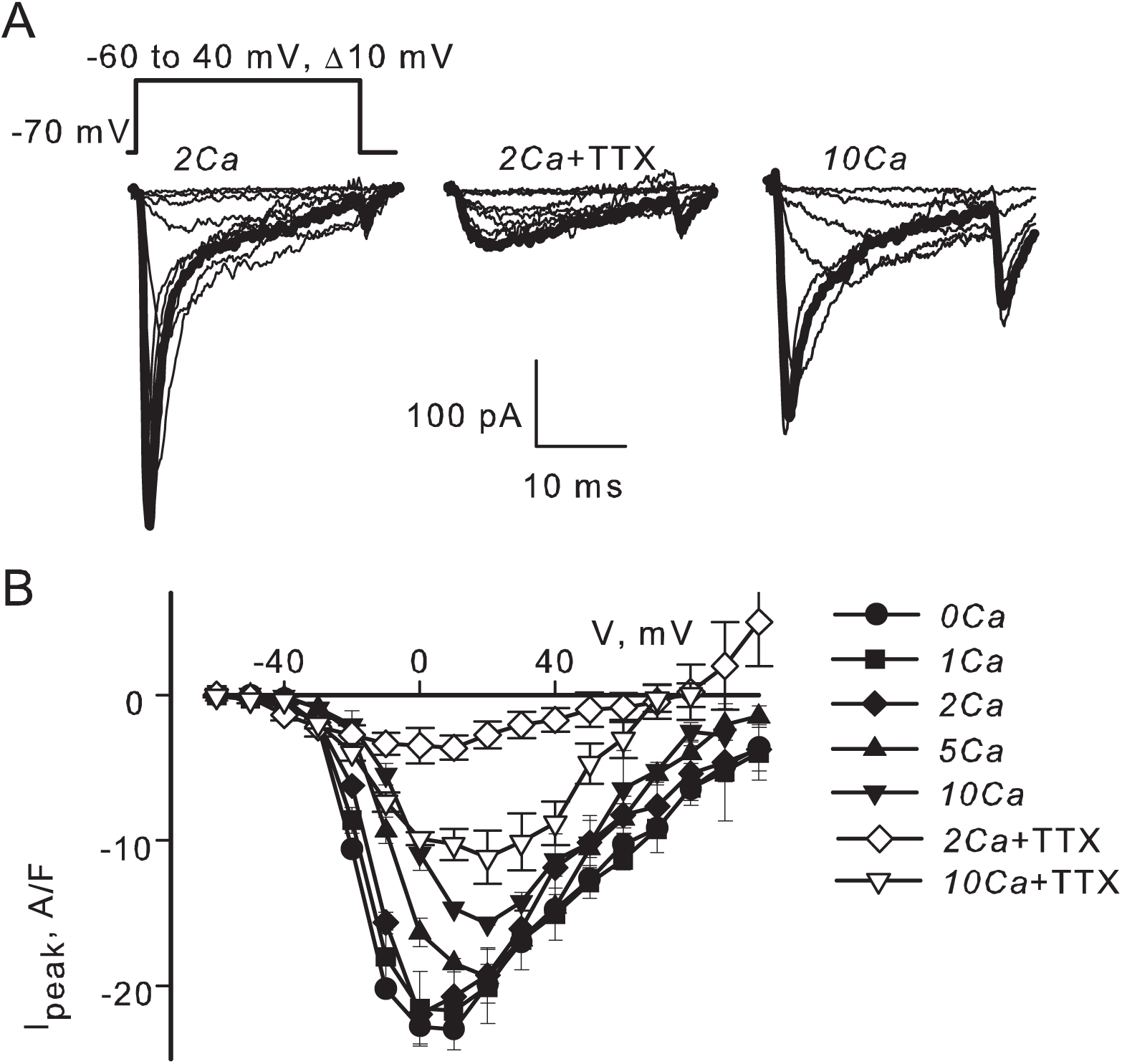
TTX-sensitive Na^+^ currents. **A)** Steps to different voltages (indicated) produced inward currents with at least two kinetically distinct components (2Ca solution). The fast component was through voltage-gated Na^+^ channels as it was blocked by application of 1 µM TTX. It was partially blocked by extracellular Ca^2+^ (10Ca solution). The thick lines highlight the maximal currents recorded at 10 mV. **B)** Averaged peak current–voltage relationships of inward currents recorded in 2Ca (n = 6) and 10Ca (n = 4) bath solutions.

Although it is expected that extracellular Ca^2+^ ions partially block TTX-sensitive Na^+^ channels [41–43], the observations in Fig 3 might also reflect an intracellular effect of Ca^2+^. Therefore, we tested the possibility that TTX-sensitive Na^+^ currents in skeletal muscle arterioles can be inhibited by elevation of Ca^2+^ on the intracellular side of the membrane. To determine whether or not intracellular Ca^2+^ regulates voltage-gated Na^+^ channels in this preparation, whole-cell currents were recorded using gramicidin-perforated voltage-clamp. Applications of 10 mM caffeine to the bath were used to reversibly increase intracellular [Ca^2+^]. Caffeine is known to have multiple effects on blood vessels [44]. Because caffeine stimulates Ca^2+^ release channels of the sarcoplasmic reticulum, it elevates intracellular bulk [Ca^2+^] well above 0.5 µM during the first 2-3 minutes of application. However, it induces only a small transient contraction in comparison to that achieved during depolarization by K^+^ (e.g., [45]). It is positioned that this happens because caffeine also accelerates Ca^2+^ re-uptake into the sarcoplasmic reticulum. The relaxing action of caffeine is attributed to its non-specific inhibition of phosphodiesterases and stimulation of cyclic nucleotide-dependent signaling pathways. In order to distinguish between the two possible modes of action by caffeine, we tested how Na^+^ currents are affected by 1 mM IBMX. IBMX bocks various phosphodiesterases with IC_50_ of 1-50 µM [46]. It has been shown that bath application of 0.5 mM IBMX reduces norepinephrine-induced vasoconstriction and elevation of intracellular [Ca^2+^] in small mesenteric arteries of rats [47].

Because the perforated patch-clamp configuration preserves the intracellular ionic content, depolarizing voltage steps produced a biphasic current illustrated in Fig 4 for steps to 0 mV. The negative peak, which gives a lower estimate of the magnitude of inward current, could be clearly distinguished at voltages between –40 and 40 mV. Both inward and outward currents were reversibly inhibited by application of 10 mM caffeine to the extracellular solution (traces a-c). The magnitude of the negative peak was reduced about two-fold. This could not result from an increase in the outward current because the outward current was nearly completely blocked. The blockage developed within 5 seconds upon caffeine application and went away in 10-15 seconds after caffeine was removed (Fig 4B, left panel). Application of 1 µM TTX to the same cell completely and reversibly blocked the negative peak, however it did not alter the outward current (Fig 4A, traces d-f). Therefore, most of the inward current was through TTX-sensitive Na^+^ channels. Averaged current-voltage relationships for the inward peak indicate that caffeine reduced TTX-sensitive currents in a voltage-independent manner (Fig 4C, n = 4). Applications of 1 mM IBMX had no measurable effect on the inward peak in four cells tested similarly (Fig 4D). Therefore, the inhibition of voltage-dependent Na^+^ currents by caffeine was likely to be due to an elevation of intracellular Ca^2+^ and not to a stimulation of the cyclic nucleotide-dependent signaling pathway.

**Fig 4.**
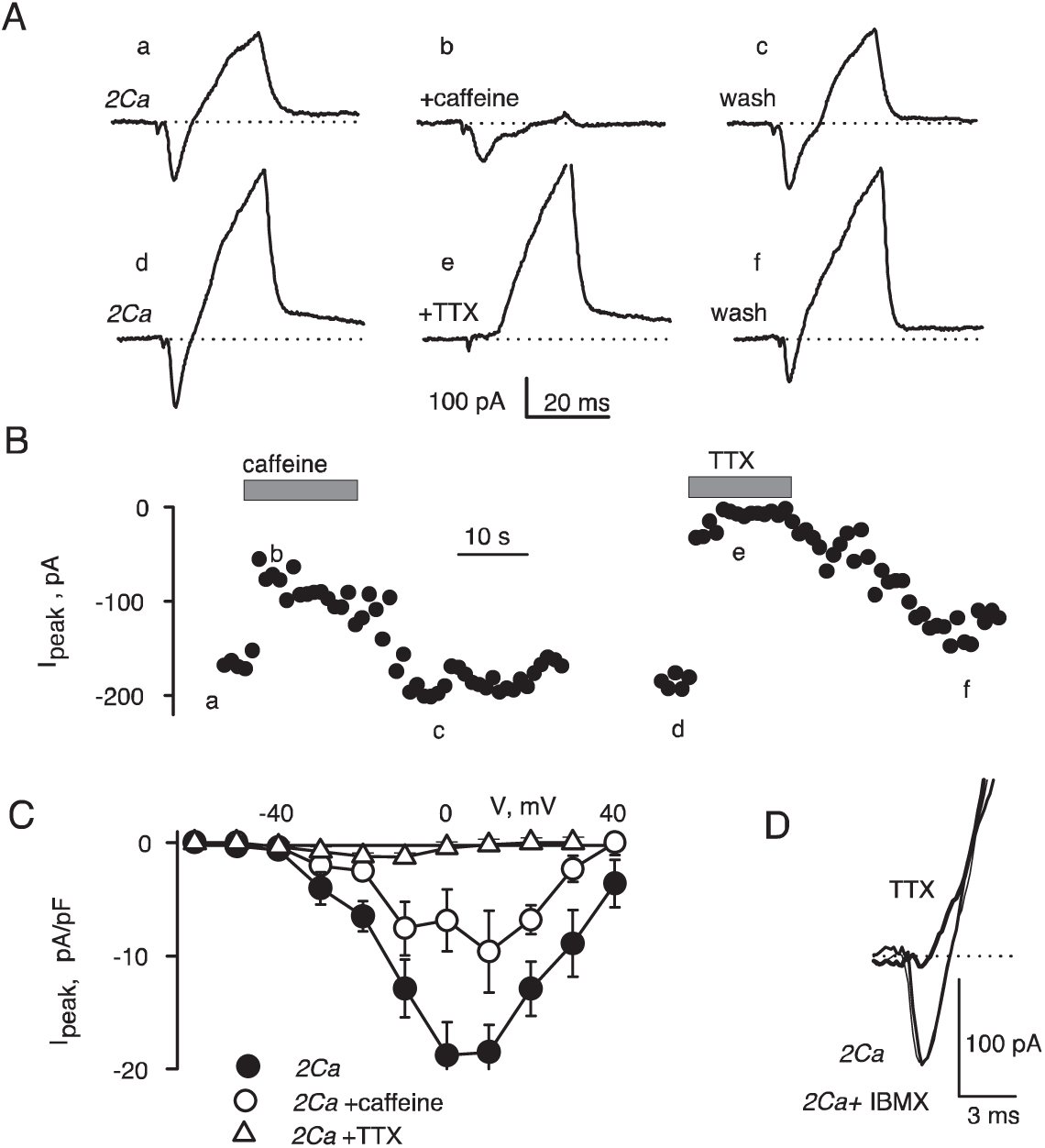
Voltage-gated Na^+^ channels depended on intracellular Ca^2+^. **A)** Whole-cell currents were recorded using gramicidin-perforated patch-clamp. Voltage steps from –70 to 0 mV produced biphasic currents with a more rapidly activating inward component (trace a, 2Ca solution). Both inward and outward currents were reduced by application of 10 mM caffeine (trace b). Effect of caffeine was reversible (trace c). Application of 1 µM TTX to the same cell completely and reversibly blocked the negative peak, but did not alter the outward current (traces d-f). **B)** Time course of blockage of the inward peak by caffeine and TTX. Time points for tracings shown in A are indicated. **C)** Averaged current-voltage relationships (n=4) for the inward peak indicate that caffeine reduced TTX-sensitive currents in a voltage-independent manner. **D)** Applications of 1 mM IBMX had no measurable effect on the inward peak in four cells tested similarly. Tracings recorded in control (2Ca) and 3 minutes after application of IBMX (indicated) overlap. However, 1 µM TTX (upper trace, indicated) completely blocked the inward peak in the same cell.

### Two types of voltage-gated Ca^2+^ channels

Because action potentials could not be completely prevented by a combined application of TTX and nifedipine (Fig 2), it is possible that a part of the depolarizing current was through nifedipine-resistant Ca^2+^ channels. To determine whether or not a part of Ca^2+^ current is through T-type channels, we characterized whole-cell Ba^2+^ currents recorded in the presence of 1µM TTX. Ba^2+^ ions dramatically reduced inactivation of L-, but not T-, type of channels. Therefore, the two kinetically different components could be more clearly observed. Steps to voltages more negative than –10 mV produced fast inactivating current (Fig 5A, panel a), and further depolarization activated slow inactivating current of greater amplitude (Fig 5A, panel b). To further isolate the more rapidly inactivating currents, we applied 10 µM of L-type Ca^2+^ channels blocker nifedipine (Fig 5B). Nifedipine blocked only the slowly inactivating component, but it did not affect the rapidly inactivating current. The nifedipine-resistant current peaked at 0 - 10 mV, which is a more positive voltage than that usually observed for T-type Ca^2+^ channels [30]. Nevertheless, the nifedipine-resistant current had several properties that are characteristic for the T-type. It inactivated somewhat faster with Ba^2+^ than with Ca^2+^ (compare traces in left and right panels in Fig 5B). Its magnitude was nearly the same in Ba^2+^ and in Ca^2+^ (Fig 5C), and it did not run-down as rapidly as the high-voltage activating nifedipine-sensitive current (data not shown).

**Fig 5.**
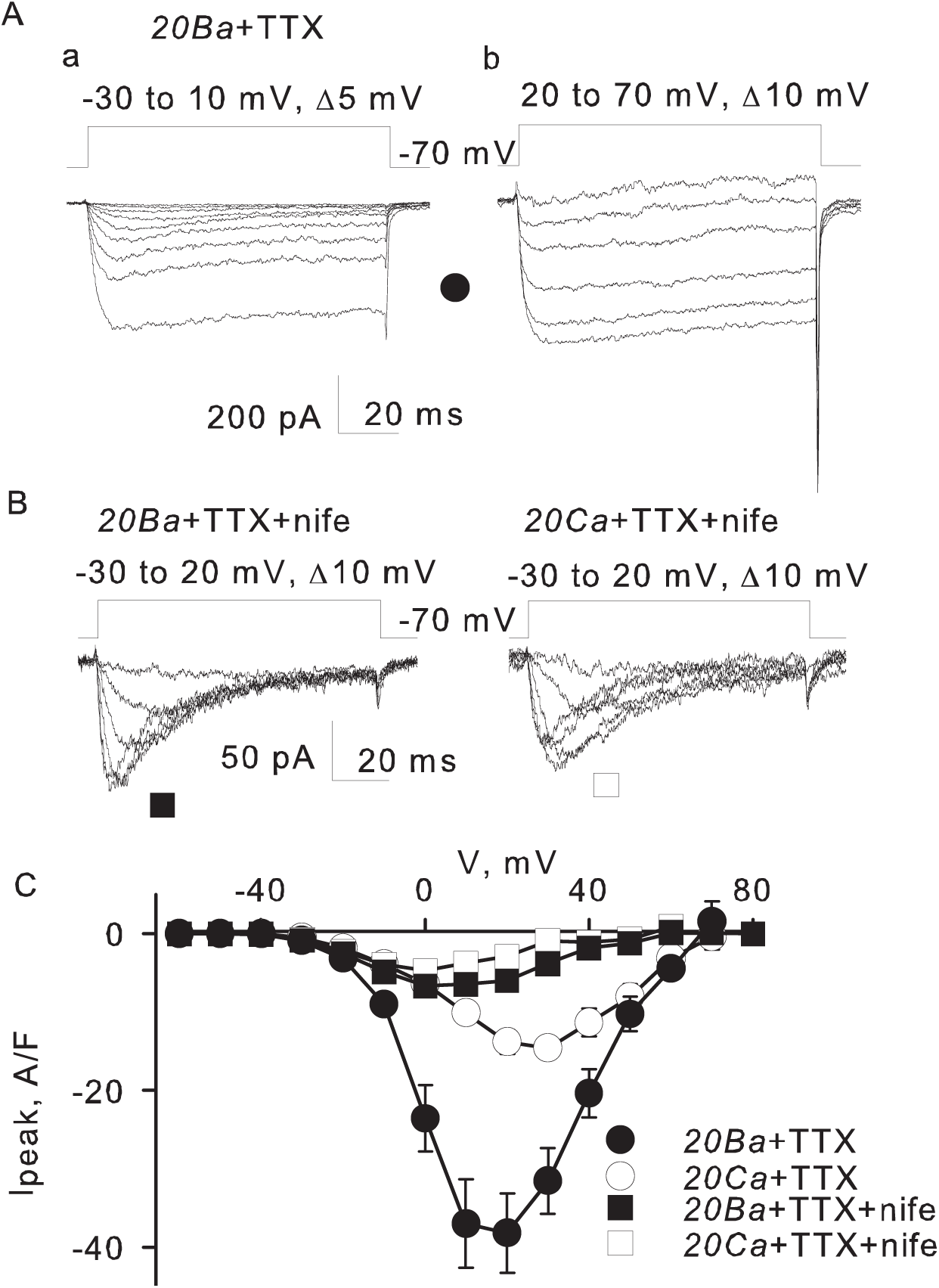
Two types of voltage-gated Ca^2+^ channels. **A)** Whole-cell Ba^2+^ currents were recorded in the presence of 1µM TTX. Two kinetically different components were observed. Voltage steps up to –10 mV produced fast inactivating current (tracings a). Further depolarization activated slow inactivating current of greater amplitude (tracings b). **B)** In the presence of 10 µM L-type Ca^2+^ channel blocker nifedipine, the rapidly activated component inactivated notably faster with Ba^2+^ (left panel) than with Ca^2+^ (right panel). **C)** Averaged peak current-voltage relationships determined in the presence of 1 µM TTX and 10 µM nifedipine as indicated. The nifedipine-resistant current peaked at about 0 mV. Its magnitude was nearly the same with Ba^2+^ and with Ca^2+^ (compare open and filled squares).

Further analysis of inactivation and deactivation kinetics supported the view that the low-voltage activating currents were through T-type Ca^2+^ channels. Using a double-pulse protocol illustrated in Fig 6, we tested effects of 100 ms pre-pulses (V1) on Ba^2+^ currents elicited by the test pulse (V2). As illustrated by the gray trace in Fig 6A (panel a), pre-pulse to –20 mV strongly reduced current at the second pulse to –20 mV. However, such a pre-pulse had little effect on current elicited by the test pulse to 30 mV (Fig 6A, panel b). The averaged dependencies of peak currents at –20 mV and 30 mV on pre-pulse voltage are shown in Fig 6B. The low- and high-voltage activating currents differ dramatically in the extent of inactivation by pre-pulses. Pre-pulses to 30 mV reduced the magnitude of low-voltage activating current by 78 ± 7 %, n = 3 (filled circles). However, similar pre-pulses decreased the high-voltage activated current only by 32 ± 8 % (open circles). The low-voltage activating currents inactivated at voltages more positive than usual for T-type Ca^2+^ channels in neurons. The voltage of half-inactivation of the low-voltage activating current was –36 ± 2 mV. The voltage-dependence of inactivation was consistent with the observation that the currents in our experimental conditions activated at more positive voltages than T-type Ca^2+^ channels in neurons.

**Fig 6.**
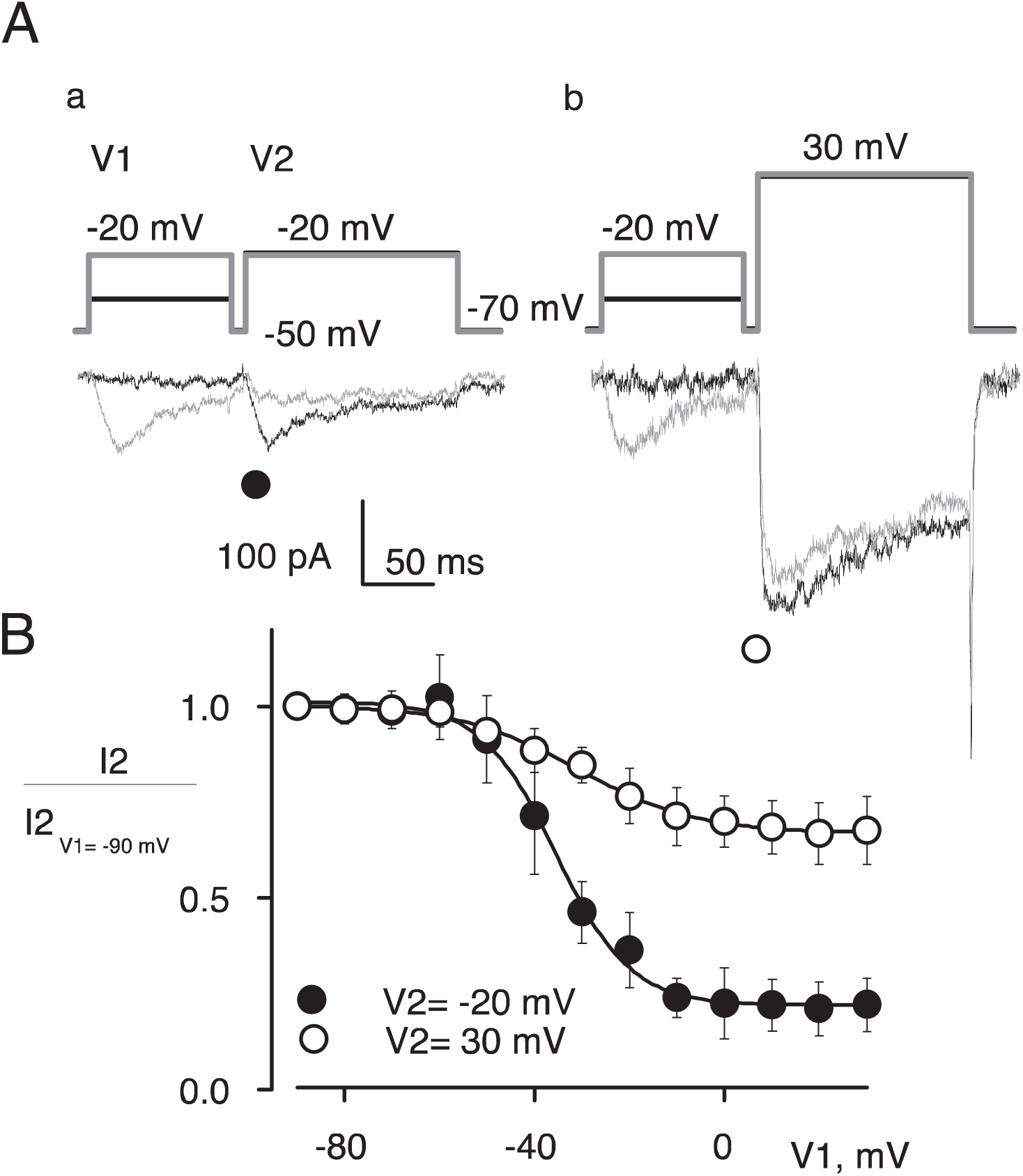
Inactivation of low-voltage activated Ca^2+^ currents. **A)** 100 ms long pre-pulse to −20 mV inactivates Ca^2+^ current that peaks at –20 mV (tracings a) but not the one that peaks at +30 mV (tracings b). **B)** Voltage-dependence of inactivation of low- (filled circles) and high- (open circles) voltage-activated Ca^2+^ currents. The smooth lines are the best fits by equation 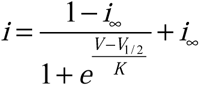

T-type channels are distinguished among other Ca^2+^ channels by their relatively slow kinetics of deactivation. To analyze low-voltage activating currents, we applied steps to different re-polarizing voltages following a 20 ms activating step to –20 mV in 20Ba+TTX solution (Fig 7A, panel a). In this case, tail currents at voltages below –20 mV had a simple mono-exponential time course. However, when the activating step was to 30 mV, the tail currents had at least two exponential components (Fig 7A, panel b). Gray lines in Fig7A illustrate the best fits at V2 = –50 mV. The averaged time constants (n = 4) of the best fits for different voltages are plotted in Fig 7B, panel a. The time constant of deactivation after stepping from –20 mV (filled squares) was 0.9 ± 0.1 ms at –120 mV and steeply increased with voltage. Parameters of the two exponential components of deactivation observed after stepping from 30 mV could be reliably determined only at voltages more positive than –90 mV. The time constant of the rapid component (open circles) was about 0.5 ms and did not change with voltage, apparently limited by the speed of the voltage clamp. At every re-polarization voltages studied, the time constant of the slower component after stepping from 30 mV (filled circles) was not significantly different from the time constant of deactivation after stepping from –20 mV (filled squares). Relative amplitudes of the deactivation components are compared in Fig 7C. Before averaging, the amplitudes of tail currents at every voltage were normalized in each cell to the amplitude of tail current elicited by stepping to –90 mV after activating pre-pulse to 30 mV. The amplitudes of the slow tail after pre-pulse to 30 mV (filled circles) and the amplitude of the tail after pre-pulse to –20 mV (filled squares) were not significantly different. Therefore, tail currents at each voltage were likely to be a sum of currents through low-voltage activating T-type Ca^2+^ channels with slow deactivation kinetics and high-voltage activating L-type Ca^2+^ channels with rapid deactivation kinetics.

**Fig 7.**
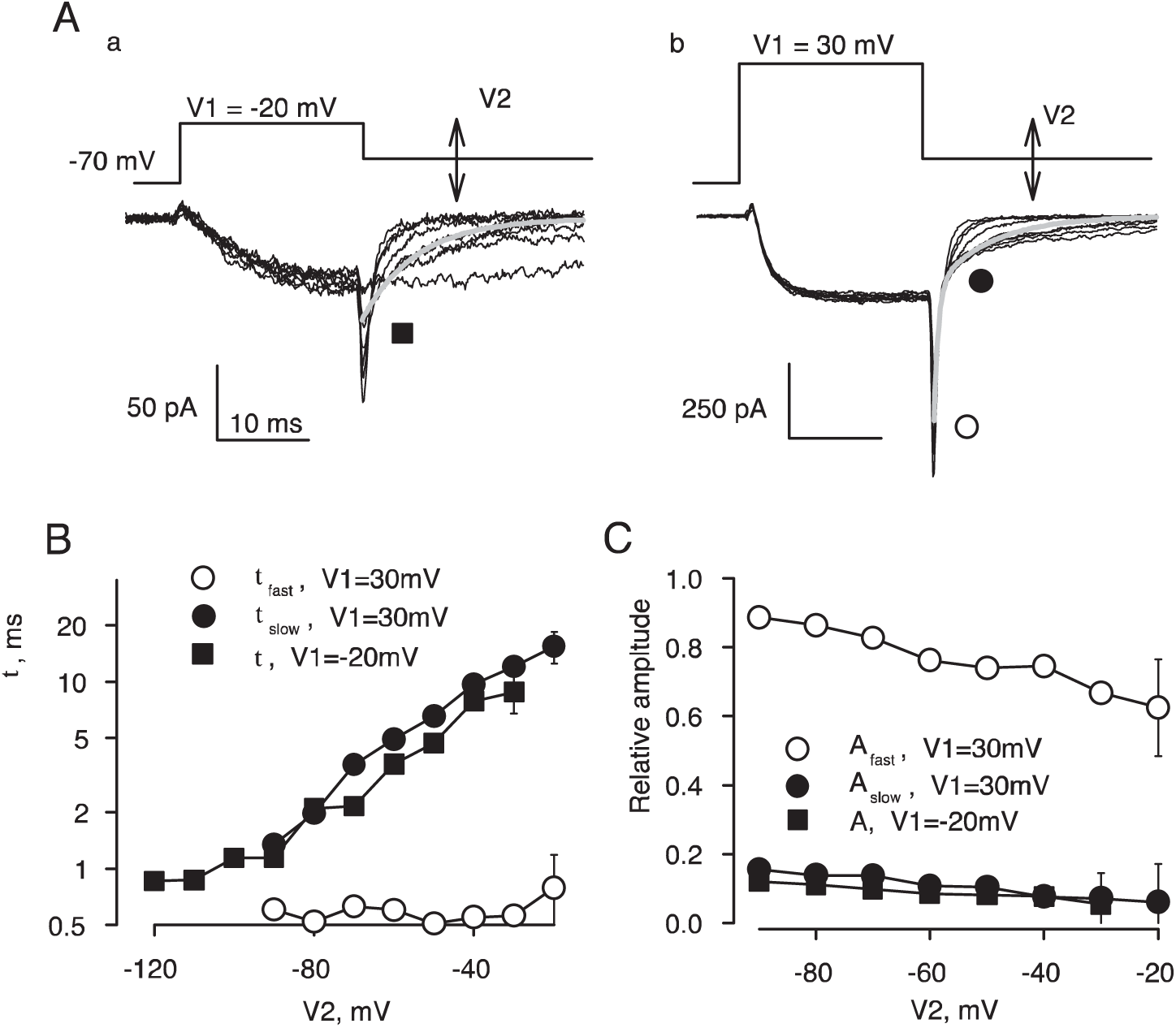
Tail currents of low-voltage Ca^2+^ channels are slow. **A)** Test voltage protocol (top) and representative tail currents for pre-pulses to −20 mV (tracings a) and +30 mV (tracings b). **B)** Averaged time constants of tail currents recorded at different voltages. **C**) The average amplitudes of tail currents recorded at different voltages. Tail currents from −20 mV were fitted by a single exponential. Tail currents from 30 mV were fitted by the sum of two exponentials. Kinetics of the slow component elicited after pre-pulse to 30 mV was similar to the tail current elicited after pre-pulse to −20 mV.

We further tested whether or not the slower component of tail currents elicited after activating pre-pulse to 30 mV passed through the low-voltage activating channels (S1 Fig). The slow component of deactivation after 30 mV pulse was nearly undetectable when a 100 ms long conditioning step to –20 mV preceded the pulse protocol shown in Fig 7A, panel b. Apparently, the low-voltage activating channels inactivated during the conditioning step and therefore did not contribute to tail currents. Unlike in the regular current-voltage relationships (e.g., Fig 5C), voltage-dependence of activation determined from the amplitude of tail currents recorded at a fixed voltage is not distorted by the changes in the electrochemical driving force and provides a more clear separation of the two components in the voltage-dependence of activation of Ba^2+^ currents (S2 Fig).

## Discussion

While Na^+^ and Ca^2+^ channels are critical for generation of action potentials in other excitable tissues, their involvement in the excitability of arteriolar smooth muscle cells is poorly understood. Both slow changes and fast spikes in the membrane potential of smooth muscle cells have been observed. The physiological role of membrane potential spikes in these cells is unclear, as relatively small changes in the membrane potential (between –55 and –35 mV) are sufficient to control Ca^2+^ entry that initiates contraction in response to stimulation by mechanical stress of the blood flow [48]. Spontaneous action potential spikes could be observed in pressurized small arteries [11, 12], as well as in un-pressurized arteries [14], and arterioles (S3 Fig). Spikes might associate with rhythmic vasomotor activity [12]; they might follow K^+^-induced hyperpolarization and dilation [23], or precede injury-induced constriction [49].

In the present study, we show that smooth muscle cells in skeletal muscle arterioles of mouse have functional voltage-gated TTX-sensitive Na^+^ and T-type Ca^2+^ channels in addition to L-type Ca^2+^ channels. All three types of channels probably contribute to the depolarizing phase of the action potential, because the rate of depolarization during the upstroke of the action potential was reduced by TTX, by nifedipine and by Ca^2+^ removal (Fig 2). The effect of TTX was most pronounced in nominally Ca^2+^-free media (Fig 2), a finding that might reflect the suppression of TTX-sensitive Na^+^ currents by elevations in cytosolic [Ca^2+^_free_] (Fig 4). Other currents also contribute to the depolarizing phase of the action potential, since depolarization was not completely suppressed by TTX in a nominally Ca^2+^-free medium. While further study is obviously needed for a better understanding of the ionic mechanisms underlying the action potential in these cells, our results clearly implicate the involvement of TTX-sensitive Na^+^ channels as well as L-type and T-type Ca^2+^ channels.

In arterioles of various vascular beds (e.g., [12, 13]), action potentials recorded with normal (1-2 mM) Ca^2+^ could appear to be Ca^2+^-dependent and insensitive to TTX because Ca^2+^ might suppress depolarizing Na^+^ currents similar to what we observed for skeletal muscle arterioles (Figs 2 and 3). Therefore, the effect of TTX in resistance arteries of other vascular beds needs to be re-examined at low Ca^2+^. Also, the molecular identity of Na^+^ channels in smooth muscle cells of skeletal muscle arterioles needs to be identified. Several isoforms of TTX-sensitive voltage-gated channels have been found previously smooth muscle cells of vasa recta (Na_V_1.3, [50]), portal vein (Na_V_1.6 and Na_V_1.8, [51]), and vas deference (Na_V_1.6, [52–54]). Our finding that caffeine, but not IBMX, reduces voltage-dependent TTX-sensitive Na^+^ currents (Fig 4) supports the view that elevations of intracellular Ca^2+^ trigger a process that suppresses activity of Na^+^ channels. Further studies are needed to investigate the exact mechanism of regulation of Na^+^ channels by Ca^2+^ in vascular smooth muscle.

TTX-sensitive Na^+^ channels are known to be controlled by Ca^2+^/calmodulin [55–60]. Ca^2+^/calmodulin binding down-regulates skeletal muscle isoform Na_V_1.4 by shifting the steady-state inactivation curve of in the hyperpolarizing direction [61]. Na_V_1.3 channels in the descending vasa recta are suppressed by calmodulin inhibitors, while elevation of the intracellular [Ca^2+^] shifts the voltage-dependence of their activation to more positive voltages [50]. Therefore, another possibility for Ca^2+^-dependent down-regulation of Na^+^ channels might be via the Protein Kinase C (PKC) pathway. It has been shown that activation of PKC decreases peak sodium currents through brain Na_V_1.2 and skeletal muscle Na_V_1.4 channels by up to 80 % [62, 63].

The exact role of T-type voltage-gated Ca^2+^ currents in normal- and pathophysiology of various vascular beds is under intense investigation. Two separate voltage-dependent Ca^2+^ currents have been found in smooth muscle cells of resistance arteries in cerebral [6, 13, 61], renal [64] and mesenteric [64–66] vascular beds. However, only L-type Ca^2+^ currents were recorded in single smooth muscle cells isolated from resistance arteries of hamster cremaster muscle [67]. Our finding of T-type Ca^2+^ currents in arterioles of leg muscles of mice is expected since both Ca_V_3.1 and Ca_V_3.2 T-type channels are expressed at the messenger RNA level in rat cremaster arterioles [32]. In our preparation, Ni^2+^ did not exhibit specificity in blockage of T-type currents. Application of 40 µM of Ni^2+^ to the bath, which should block primarily Ca_V_3.2 channels [30], had only partial effect (46 ± 18 % reduction, n=5, data not shown). Therefore, we suggest that both Ca_V_3.1 and Ca_V_3.2 channels were present. It is not clear why T-type Ca^2+^ currents were not found in hamster cremaster [67]. It is possible that the critical difference between our preparation in this study and that of Cohen and Jackson [67] is that we recorded from smooth muscle cells that were situated in the vessel and not separated into single cells. Although the in-vessel recordings are disadvantageous for the whole-cell patch-clamp configuration because of the electrical coupling between smooth muscle cells, the overall condition of the cells is much better and closer to their physiological state.

Several groups have proposed that Ca^2+^ influx through T-type Ca^2+^ channels contributes to vascular contractility [32, 68]. However, mice with knocked-out gene of the Ca_V_3.2 T-type channel die from a constitutive spasm of coronary arteries because of impaired relaxation of smooth muscle cells [33, 69]. Although no direct measurements of membrane potential were performed, this finding strongly indicates that the T-type Ca^2+^ channel is essential for relaxation of smooth muscle cells.

Localized changes of sub-membrane Ca^2+^ due to constitutively active L-type Ca^2+^ channels (sparklets) have been proposed to be primary for both the near-membrane and global Ca^2+^ control of smooth muscle cell contractility [70, 71]. The voltage-dependence of sustained elevation of the bulk intracellular Ca^2+^ due to L-type Ca^2+^ channels was found to be from –40 to –15 mV, peaking at –30 mV [72]. In neurons, the voltage range for sustained activity of T-type Ca^2+^ channels is from –90 to –30 mV [73]. Although we found that T-type Ca^2+^ channels in smooth muscle cells of skeletal muscle arterioles activate at least 20 mV more positively than in neurons, their activation occurs at more negative voltages in comparison with L-type Ca^2+^ channels (S2 Fig). Relevant to the voltage-dependence of activation of T-type Ca^2+^ channels, we found that the resting membrane potential in our experimental conditions was about –70 mV (Fig 1). Similar resting potentials were previously reported for un-pressurized mesenteric artery [74], submucosal arterioles [75], cerebral arterioles [13, 76]. However, it is significantly more negative than that in un-pressurized arterioles in rat cremaster muscle (about –55 mV, [12]), which could be due to the difference between gramicidin-perforated patch clamp and conventional microelectrode techniques. Gramicidin-perforated patch clamp is concievably less damaging for small cells as it prevents ionic leaks through the giga-seal and does not alter intracellular [Cl^−^] [77]. Another reason for more negative resting membrane potential could be that our solutions didn’t include HCO_3_^−^. The HCO_3_^−^Cl^−^ exchanger in the plasma membrane of arteriolar smooth muscle hs been shown to contribute to the high intracellular [Cl^−^] [78].

## Conclusion

To summarize, we developed a methodology that allows for the isolation and study of murine skeletal muscle arterioles in ex-vivo preparation. We confirmed that smooth muscle cells of skeletal muscle arterioles express L-type voltage-gated Ca^2+^ channels as well as TTX-sensitive Na^+^ and T-type Ca^2+^ channels. More specifically, we recorded voltage-dependent inward currents using voltage- and current-clamp recordings, including perforated whole-cell path-clamp technique. We also examined potential contributions of these channels to the excitability of smooth muscle cells by electrophysiology and pharmacology. Our data demonstrate two types of voltage-gated Ca^2+^ channels and TTX-sensitive Na^+^ channels respond to depolarization of smooth muscle cells. Their physiological properties and, hence, relative contributions may change depending on the state of the arteiolar vasculature. T-type Ca^2+^ channels can be tuned both for depolarization (constriction) and Ca^2+^-dependent stabilization of negative voltages (dilation, via activation of BK_Ca_ channels) [79]. Ca^2+^ influx through the low-voltage activated T-type Ca^2+^ channels might serve to inhibit Na^+^ channels and thus prevent the occurrence of action potential spikes.

## Acknowledgements

The authors are grateful to John Reeves (Rutgers University), Sergey Smirnov (University of Bath), and Natalia Shirokova (Rutgers University) for valuable suggestions and discussions.

## Supporting information

**S1Fig.**
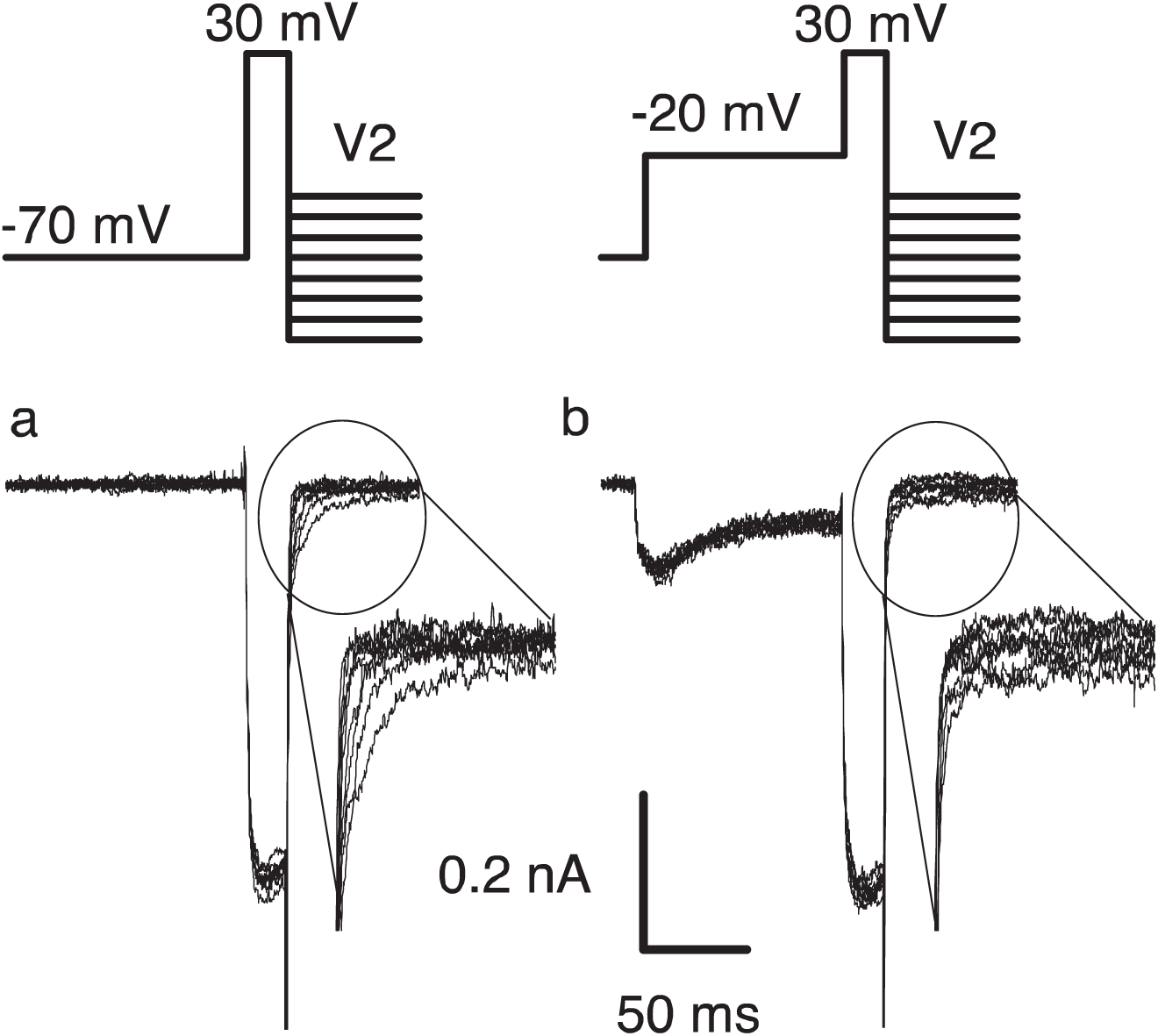
The effect of conditioning on slow tail currents. The tail currents elicited after activating step to 30 mV had significant slow component (tracings a). The slow component was absent if the conditioning that inactivates currents at –20 mV preceded the activating step (tracings b). Similar observations were done on three cells.

**S2 Fig.**
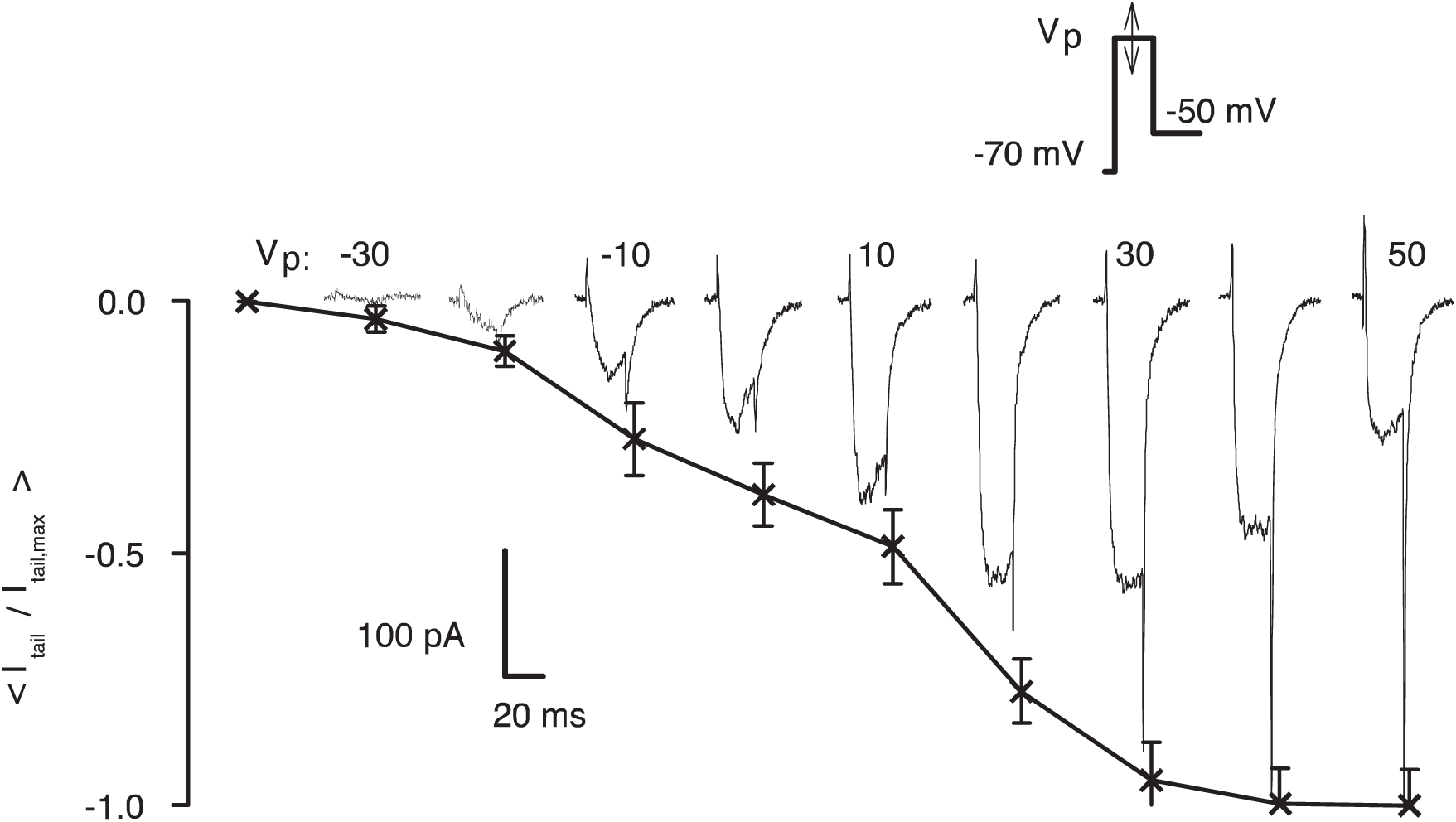
Magnitude of tail current reveals activation of Ca^2^+^^ channels with different voltage-dependence. Test voltage protocol is shown at the top. Traces of representative currents for different pre-pulse are shifted along the *x-*axis by intervals proportional to the pre-pulse voltage (indicated). Before averaging (n = 3), amplitudes of the tail current in each cell were normalized to their value for activation step to 50 mV.

**S3 Fig.**
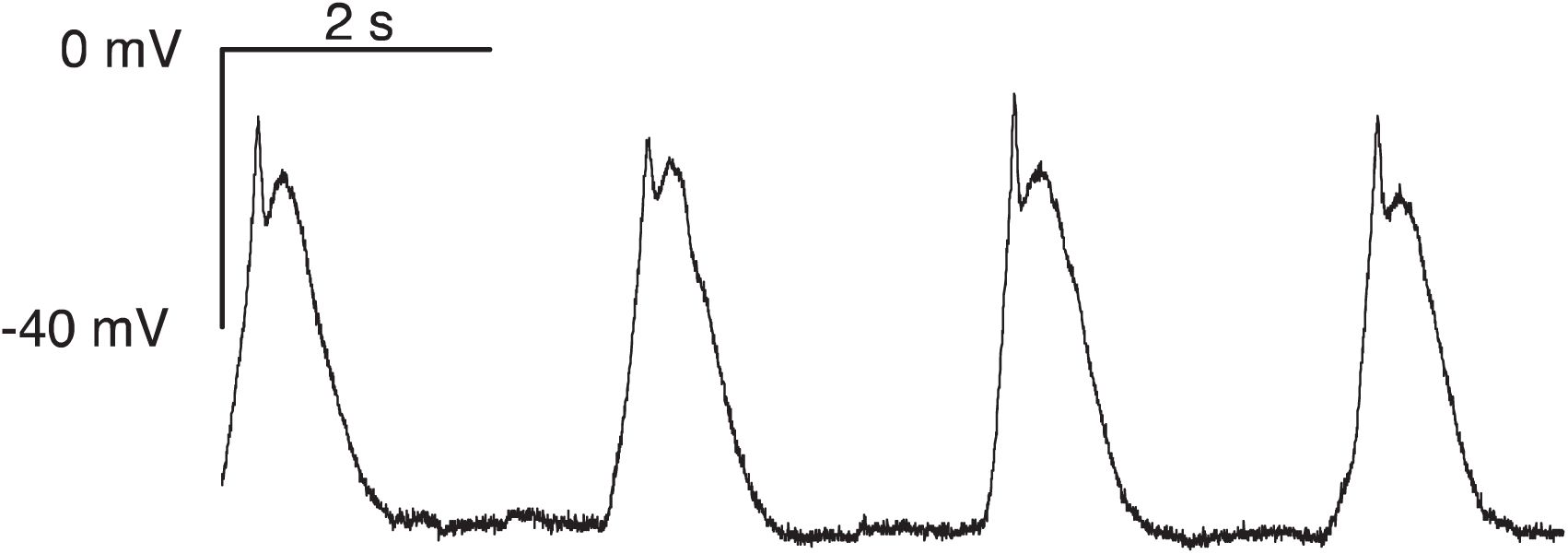
An example of spontaneous action potentials in smooth muscle cells of skeletal muscle arterioles. Recording conditions: gramicidin-perforated patch-clamp with 2Ca bathing solution and 150 mM KCl in the pipette. Similar rhythmic activity was observed in six out of 81 cells studied. Most cells had stable membrane potential.

